# Emergence of an Australian-like *pstS*-null vancomycin resistant *Enterococcus faecium* clone in Scotland

**DOI:** 10.1101/236786

**Authors:** Kimon Lemonidis, Talal S. Salih, Stephanie J. Dancer, Iain S. Hunter, Nicholas P. Tucker

## Abstract

Multi-locus sequencing typing (MLST) is widely used to monitor the phylogeny of microbial outbreaks. However, several strains of vancomycin-resistant *Enterococcus faecium* (VREfm) with a missing MLST locus (*pstS*) have recently emerged in Australia, with a few cases also reported in England. Here, we identified similarly distinct strains circulating in two closely located hospitals in Scotland. Whole genome sequencing of five VREfm strains isolated from these hospitals identified four *pstS*-null strains across both hospitals, while the fifth was of a multi-locus sequence type (ST) 262, which is the first documented in the UK. All five Scottish isolates had an insertion in the *tetM* gene, which is associated with increased susceptibility to tetracyclines, providing no other tetracycline-resistant gene is present. Such an insertion, which encompasses a *dfrG* gene and two currently uncharacterised genes, was additionally identified in all tested *VanA*-type *pstS*-null VREfm strains (5 English and 18 Australian). Phylogenetic comparison with other VREfm genomes indicates that the four *pstS*-null Scottish isolates sequenced in this study are more closely related to *pstS*-null strains from Australia rather than the English *pstS*-null isolates. Given how rapidly such *pstS*-null strains have expanded in Australia, the emergence of this clone in Scotland raises concerns for a potential outbreak.

## Introduction

Vancomycin-resistant *Enterococcus* (VRE) was first identified about three decades ago and has now become a major nosocomial pathogen. It typically infects immunocompromised patients and can cause endocarditis, bloodstream, urinary tract, and skin and skin structure infections [1]. VRE infections are generally more serious than those caused by vancomycin-susceptible enterococci, and are associated with higher mortality rates [2]. Among all VRE species, vancomycin-resistant *Enterococcus faecium* (VREfm) is responsible for the majority of hospital infections. VREfm has been recently listed as a high priority pathogen for research and development of new antibiotics by the World Health Organisation [3]. Various measures have been implemented to monitor the spread of VREfm infections, including multi-locus sequence typing (MLST); this relies on characterising the allelic profile of seven “house-keeping” genes, located in the *E. faecium* chromosome [4]. Although useful, this MLST scheme is of limited resolution to accurately capture the clonal type of isolated *E. faecium* strains [5–7]. Furthermore, VREfm isolates lacking the *pstS* MLST-gene locus have recently emerged in both England [7] and Australia [8].

Here, we have used whole genome sequencing to identify four VREfm isolates from two Scottish hospitals, which lack the *pstS* locus, and another Scottish VREfm isolate with a MLST profile that, to the best of our knowledge, has not been previously reported in the UK. Additionally, we provide information regarding their resistance profiles and epidemiology.

## Materials and Methods

VRE strains were isolated from 5 patients in 2 Scottish hospitals, between January to October 2017; these patients developed complications following either biliary or colonic surgery, and had been treated with various combinations of penicillin, amoxicillin, flucloxacillin, Tazocin™, gentamicin, metronidazole and vancomycin during their hospital stay. The VRE strains were cultured on horse blood agar with single colonies transferred to Mueller Hinton broth (Oxoid) liquid. These were subsequently placed on Mueller Hinton agar; single colony cultures of these were subsequently used for DNA isolation. Genomic DNA was extracted using an Isolate II genomic DNA kit (Bioline), using the manufacturer’s instructions for difficult to lyse Gram-positive bacteria. DNA libraries were then prepared using the NEBNext^®^ Fast DNA Fragmentation and Library Prep Set for Ion Torrent™ (New England Biolabs): briefly, 1 μg of plasmid DNA was fragmented, and Ion Xpess barcode adapters (Life Technologies) were ligated to the DNA fragments; after clean-up using Agencourt AMPure XP beads (Beckman Coulter), 400 bp target fragments were isolated following 18 min electrophoresis on E-gel^®^ SizeSelect™ agarose gels (Life Technologies); these were subsequently amplified by PCR and, following another clean-up with Agencourt^®^ AMPure^®^ XP beads, the quality of the resulting DNA libraries was assessed on a 2100 Bioanalyzer^®^ (Agilent Technologies), using high sensitivity DNA chips (Agilent Technologies). Template positive Ion Sphere™ particles (ISPs) for semiconductor sequencing were prepared using the Ion Touch 2 System (Life Technologies). Enriched ISPs were loaded into ion v2 BC 316™ chips (2 genomes per chip) and sequenced on an Ion PGM™ system (Life Technologies). Low quality reads (quality score threshold: 0.05) were trimmed using CLC genomics Workbench (Qiagen, version 9.5.2), and resulting reads were assembled using SPAdes (St. Petersburg genome assembler, version 3.9). Contigs having less than 1000 bp sequences were discarded. The remaining contigs were reordered on Mauve (version 20150226) using the *E. faecium* Aus0004 genome [9] as reference, and resulting genome sequences were submitted to GenBank under the assessions PJZU00000000 (VREF001), PJZT00000000 (VREF002), PJZS00000000 (VREF003), PJZR00000000 (VREF004) and PJZQ00000000 (VREF005). Antibiotic resistant genes were identified as perfect, strict or loose matches against resistant genes of the comprehensive antibiotic resistance database (CARD) [10]. Sequence reads of *pstS*-null genomes from England and Australia were obtained from European Nucleotide Archive, and reads were assembled using CLC genomics Workbench; contigs having less than 200 bp were discarded. *In silico* MLST analysis was performed using the PubMLST website (http://pubmlst.org/) [11]. Alignments of genomes were done using the REALPHY (Reference sequence Alignment based Phylogeny builder) online tool (version 1.12) [12], with *E. faecium* Aus0004 [9], Aus0085 [13] as references. Minimum inhibitory concentrations (MICs) for vancomycin, streptomycin, spectinomycin, tetracycline, oxytetracycline, doxycycline, minocycline and rifampicin were calculated using the microdilution method in cation adjusted Mueller-Hinton broth media (BD Biosciences), whereas MICs for all other antibiotics were obtained using VITEK 2 (bioMerieux).

## Results

All Scottish VRE strains sequenced (VREF001-5) had 2.9-3.0 Mb genomes with 37.6-37.7% GC content (Table 1). Analysis of average nucleotide identity and Tetra Correlation Search (TCS) against database genome sequences, using JSpeciesWS (version 3.0.12) [14], showed that all five VRE isolates were *E. faecium;* VREF001, VREF002, VREF004 and VREF005 were highly related with over 99.9% identity among aligned (>97%) sequences, whereas VREF003 displayed around 16% unique sequences compared with the other 4 genomes. *In silico* MLST analysis of VREF001-5 genomes identified VREF003 as sequence type (ST) 262, which has not been previously reported in the United Kingdom. The other 4 genomes had exact matches for 6 MLST alleles (atpA-9, ddl-1, gdh-1, *purK-1*, gyd-12, *adk-* 1), but no match for *pstS*; such MLST profile has now been assigned as ST1424 in the *Enterococcus faecium* MLST database (https://pubmlst.org/efaecium/). However, despite the missing *pstS* in these four genomes, all VREfm strains sequenced in this study did have a *pstS* homologue within a *pst* operon (also referred to as *pstS2*), which is thought to be the actual *pstS* housekeeping gene in *E. faecium* [7,8].

**Table 1.**
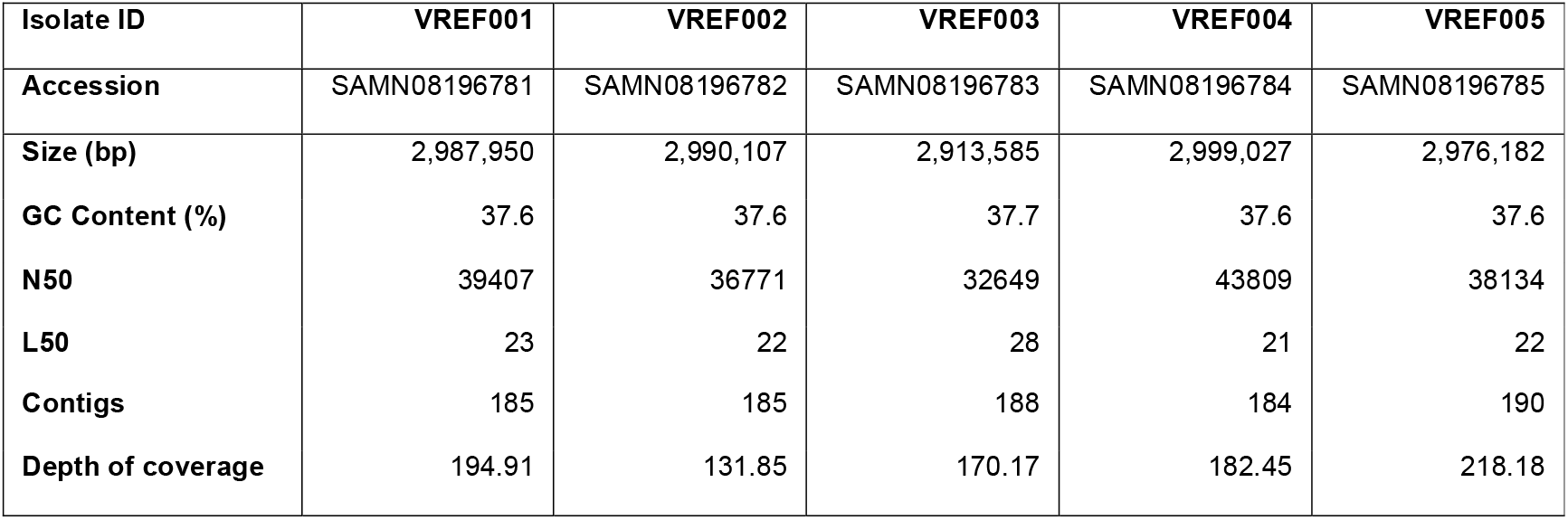
Features of assembled sequences of the five isolates.

| Isolate ID | VREF001 | VREF002 | VREF003 | VREF004 | VREF005 |
| --- | --- | --- | --- | --- | --- |
| Accession | SAMN08196781 | SAMN08196782 | SAMN08196783 | SAMN08196784 | SAMN08196785 |
| Size (bp) | 2,987,950 | 2,990,107 | 2,913,585 | 2,999,027 | 2,976,182 |
| GC Content (%) | 37.6 | 37.6 | 37.7 | 37.6 | 37.6 |
| N50 | 39407 | 36771 | 32649 | 43809 | 38134 |
| L50 | 23 | 22 | 28 | 21 | 22 |
| Contigs | 185 | 185 | 188 | 184 | 190 |
| Depth of coverage | 194.91 | 131.85 | 170.17 | 182.45 | 218.18 |

We then searched for antibiotic resistance genes with perfect or strict matches to reference genes of the comprehensive antibiotic resistance database (CARD), but also considering loose matches with exceptional low e-value (<10^-100^) and/or very high identity (>66%) to the reference gene. Analysis of the gene profiles of these five isolates identified the multi-aminoglycoside resistance gene *aac(6′)-Ii*, in all 5 strains, and *aac(6′)-aph(2″*), in all but VREF004; all 5 isolates had at least 1 macrolide-lincosamide-streptogramin resistance gene (*ermB*), a *pbp5* variant (designated *pbp5-R*) conferring resistance to beta-lactams [15], for which no reference gene exists in CARD, and the vancomycin-teicoplanin resistance gene, *vanA*. With the exception of VREF003, all strains had the spectinomycin resistance gene *ant(9)-Ia*. Lastly, VREF003 was the only strain with a tetracycline transporter gene (*tetL*) (Table 2). It is notable that all isolates had 2 loose matches for the tetracycline resistance gene *tetM*. By further examining the region where these *tetM*-like sequences are located, we found that the 5 genomes had a Tn5801-like transposon (full-length or fragment) containing a sequence similar to *Staphylococcus aureus* Mu50 *tetM* gene [16], but with a 3,229 bp insertion. The trimethoprim-resistant gene *dfrG* and two more genes with unknown function were located within this insertion. A similar insertion has been reported previously in some ST17 and ST18 *E. faecium* strains [16] which is thought to result in defective protection against tetracyclines [17]. Lastly, all strains had a homologue of the *E. faecalis IsaA* gene which confers resistance to quinipristin-dalfopristin. However, the product of this gene (known as EatA), unlike the >99% identical EatA_v_ product found in some *E. faecium* strains, has a threonine at position 450, which is associated with increased susceptibility towards quinupristin-dalfopristin [18,19].

**Table 2.** MIC values (mg/L) of selected antibiotics against the 5 Scottish VREfm isolates. Susceptibility to a given antibiotic is indicated in bold. ND: MIC value not determined. Resistant genes identified which correspond to strict or perfect matches against antibiotic resistance genes of the comprehensive antibiotic resistance database (CARD) are shown. Additionally, some loose matches (underlined) to CARD reference genes, which either had > 99% identity with reference CARD gene (*tetL*), or had zero e-value (*rpoB2*), or were the sole genes that could explain differences in antimicrobial susceptibility among the 5 isolates (*ant(9)-Ia, marA* and *mtrR*), are shown. The variant of a full-length penicillin-binding protein 5 gene conferring resistance to penicillins (*pbp5-R*), for which no reference gene in CARD exists, was found in all genomes. Numbers at top right of resistant genes correspond to the sequenced strains (last digit of isolate ID) carrying these genes.

|  |  | VREF00<br>1 | VREF00<br>2 | VREF00<br>3 | VREF00<br>4 | VREF00<br>5 | Resistance genes |
| --- | --- | --- | --- | --- | --- | --- | --- |
| **Beta-lactams** | Amoxicillin | > 32 | > 32 | > 32 | > 32 | > 32 | *pbp5-R*<sup>1-5</sup> |
|  | Co-Amoxiclav | > 8 | > 8 | > 8 | > 8 | > 8 | *pbp5-R*<sup>1-5</sup> |
| **Glycopeptides** | Vancomycin | 64 | 512 | > 512 | 512 | 128 | *vanA*<sup>1-5</sup>, *vanHA*<sup>1-3,5</sup>, *vanRA*<sup>1-5</sup>, *vanSA*<sup>1-4</sup>,<br>*vanXA*<sup>1-5</sup>, *vanYA*<sup>2-4</sup>, *vanZA*<sup>3</sup> |
|  | Teicoplanin | > 32 | > 32 | > 32 | > 32 | 4 | *vanA*<sup>1-5</sup>, *vanHA*<sup>1-3,5</sup>, *vanRA*<sup>1-5</sup>, *vanSA*<sup>1-4</sup>,<br>*vanXA*<sup>1-5</sup>, *vanYA*<sup>2-4</sup>, *vanZA*<sup>3</sup> |
| **Fluoroquinolones** | Ciprofloxacin | > 4 | > 4 | > 4 | > 4 | ND | *efmA*<sup>1-5</sup> |
| **Aminoglycosides** | Gentamicin | > 128 | > 128 | > 128 | > 128 | > 128 | *aac(6')-aph(2'')*<sup>1-3,5</sup> |
|  | Kanamycin | > 128 | > 128 | > 128 | > 128 | > 128 | *aac(6')-aph(2'')*<sup>1-3,5</sup>, *aph(3')-IIIa*<sup>1-2,4-5</sup>,<br>*aac(6')-li*<sup>1-5</sup> |
|  | Streptomycin | 32 | 32 | 32 | 32 | 32 | None |
|  | Spectinomycin | 512 | 512 | 32 | > 512 | 512 | *ant(9)-la*<sup>1-2,4-5</sup> |
| **Tetracyclines** | Tetracycline | ≤ 0.25 | ≤ 0.25 | > 128 | ≤ 0.25 | ≤ 0.25 | *tetL*<sup>3</sup> |
|  | Oxytetracycline | 1 | ≤ 0.25 | > 128 | 0.5 | ≤ 0.25 | *tetL*<sup>3</sup> |
|  | Doxycycline | ≤ 0.25 | ≤ 0.25 | 16 | ≤ 0.25 | ≤ 0.25 | *tetL*<sup>3</sup> |
|  | Minocycline | ≤ 0.13 | ≤ 0.13 | 16 | ≤ 0.13 | ≤ 0.13 | *tetL*<sup>3</sup> |
|  | Tigecycline | ≤ 0.13 | ≤ 0.13 | ≤ 0.13 | ≤ 0.13 | ≤ 0.13 | None |
| **Macrolides,**<br>**Lincosamides,**<br>**Streptogramins** | Clarithromycin | > 4 | > 4 | > 4 | > 4 | > 4 | *ermA*<sup>1-2,4-5</sup>, *ermB*<sup>1-5</sup> |
|  | Quinupristin-Dalfopristin | 0.5 | 0.5 | 0.5 | 1 | 0.5 | None |
| **Others** | Chloramphenicol | 8 | 8 | ≤ 4 | 8 | 8 | None |
|  | Linezolid | 2 | 2 | 2 | 2 | 2 | None |
|  | Nitrofurantoin | 128 | 256 | 128 | 256 | 256 | Unknown |
|  | Trimethoprim | > 16 | > 16 | > 16 | > 16 | > 16 | *dfgG*<sup>1-5</sup> |
|  | Rifampicin | 16 | 16 | 1 | 16 | 16 | *rpoB2*<sup>1-5</sup>, *marA*<sup>1-2,4-5</sup>, *mtrR*<sup>1-2,4-5</sup> |

All strains exhibited high level resistance to vancomycin; 4/5 had high level resistance to teicoplanin, associated with the presence of *vanSA* (absent from VREF005, which displayed low level resistance) and 4/5 had high level resistance to spectinomycin (apart from VREF003 which displayed low level resistance). The strains had low level resistance to streptomycin, as well as resistance to a variety of other drugs. The low level resistance to streptomycin and spectinomycin, in the absence of any gene coding for resistance to those antibiotics, is due to decreased uptake of these drugs by *E. faecium* [20]. All isolates were susceptible to linezolid, chloramphenicol, tigecycline and Synercid™ (quinupristin-dalfopristin). VREF003 was the only isolate found to be resistant to tetracycline, doxycycline, oxytetracycline and minocycline, but unlike the other four isolates, it was susceptible to rifampicin (Table 2). Since all five isolates had an identical *rpoB2* variant (assumed to exert some level of rifampicin-resistance), the decreased resistance of VREF003 towards rifampicin, compared to the other 4 strains, may be due to the lack of any *mtrR* and *marA* homologues in the VREF003 genome. On the other hand, loose matches of these were present in the other four isolates, and such genes have been shown to confer resistance to multiple drugs in other bacteria, including rifampicin [21,22]. Alternatively, VREF001-2 and VREF004-5 may contain an *mtrR*-independent mechanism of reduced permeability for rifampicin compared with VREF003 [23]. Antibiotic susceptibility tests indicated that all five isolates are classified as multi-drug resistant enterococci, according to standardised international terminology [24].

We then performed a phylogenetic comparison of VREF001-5 genomes with other known VREfm genomes reported to have a missing *pstS* locus; these included: 14 (out of 66 reported) Australian strains isolated from 9 different hospitals between 2014 and 2015 [8], 5 Australian strains (out of 202 reported within 6 local health districts) isolated from 2 different local health districts (LHD-1 and LHD-2) in New South Wales during 2016 [25], and all 5 English *pstS*-null *E. faecium* strains isolated from a single hospital (Kathy Raven, personal communication) in 2005 [7]. In our phylogenetic analysis we also included: 5 hospital-associated isolates of different MLST sequence type each, 3 isolates (BM4538, UAA1025 and E2883) previously reported to have an insertion in the *tetM*-locus [16] and the first complete *E. faecium* genome, Aus0004 [9].

This analysis indicated that Scottish *pstS*-null isolates are most closely related to Australian *pstS*-null isolates (Fig. 1A; subclade 3-1), and particularly to certain ST1424 clones isolated from New South Wales (hospital 3 and Local Health District 1). Due to the existence of long branches within subclade 3-1 (Fig. 1A), the phylogeny of that subclade was further refined by analysis of genomes belonging only to this subclade [12] (Fig. 1B); most Australian ST1421 clones and the sole ST1422 were found to be closely related to the ST1424 subclade; however, few Australian ST1421 strains (SVH-244, DMG1500788 and DMG1500808) and one ST1424 (SVH-278) were more phylogenetically distinct, whereas the sole ST1423 clone (*VanB* type DMG1500761) was found to diverge the most among all *pstS*-null strains (Fig. 1A-C). English *pstS*-null isolates, which are all ST1477, all cluster together (Fig. 1A; subclade 2-2) and are distinct form Australian and Scottish *pstS*-null strains; nevertheless, all *pstS*-null clones appear to be related to ST17 clones (Fig. 1A; subclade 3-2 and Fig. 1C), which suggests that the former may have derived from ST17 strains. Most of the Australian *pstS*-null genomes we analysed were ST1421 (Fig. 1C), in agreement with recent data showing that ST1421 is the most common and widespread *pstS*-null VRE strain in Australia, accounting for more than 70% of the cases [25,26]. Since both Australian [8] and Scottish (this study) VREfm *pstS*-null strains were found in different hospitals (Fig. 1C), it is likely that such clones have spread in the community. Our phylogenetic analysis further supports an intercontinental ST1424 clone spread between hospitals of New South Wales and Scotland (Fig. 1). On the other hand, all English VREfm strains with missing *pstS* locus [7] cluster together and are likely to have spread within the hospital that they were isolated from. VREF003 is more phylogenetically related to ST18 *E. faecium* isolates (Fig. 1A), in agreement with a previous study showing a phylogenetic relationship between ST262 and ST18 *E. faecium* strains [27]. *VanA* type VREfm strains lacking the *pstS* locus (4/4 Scottish, 18/18 Australian and 5/5 English) were found to also contain a *Tn5801-like* transposon with an insertion in the *tetM* locus (Fig. 1C). This genotype, previously documented in the MLST-typeable Aus0004, BM4538, E2883 and UAA1025 [16] included in this analysis, was further observed in VREF003, ERR374934 and ERR374968. The sole *vanB* type VREfm with missing *pstS* reported, though, had an intact *tetM* gene sequence. Collectively, it appears that the *tetM* insertion in VREfm is associated with *VanA* type strains missing *pstS* (STs: 1421, 1422, 1424 and 1477), as well as with STs: 17, 18, 80 and 262 (Fig. 1C).

**Figure 1.**
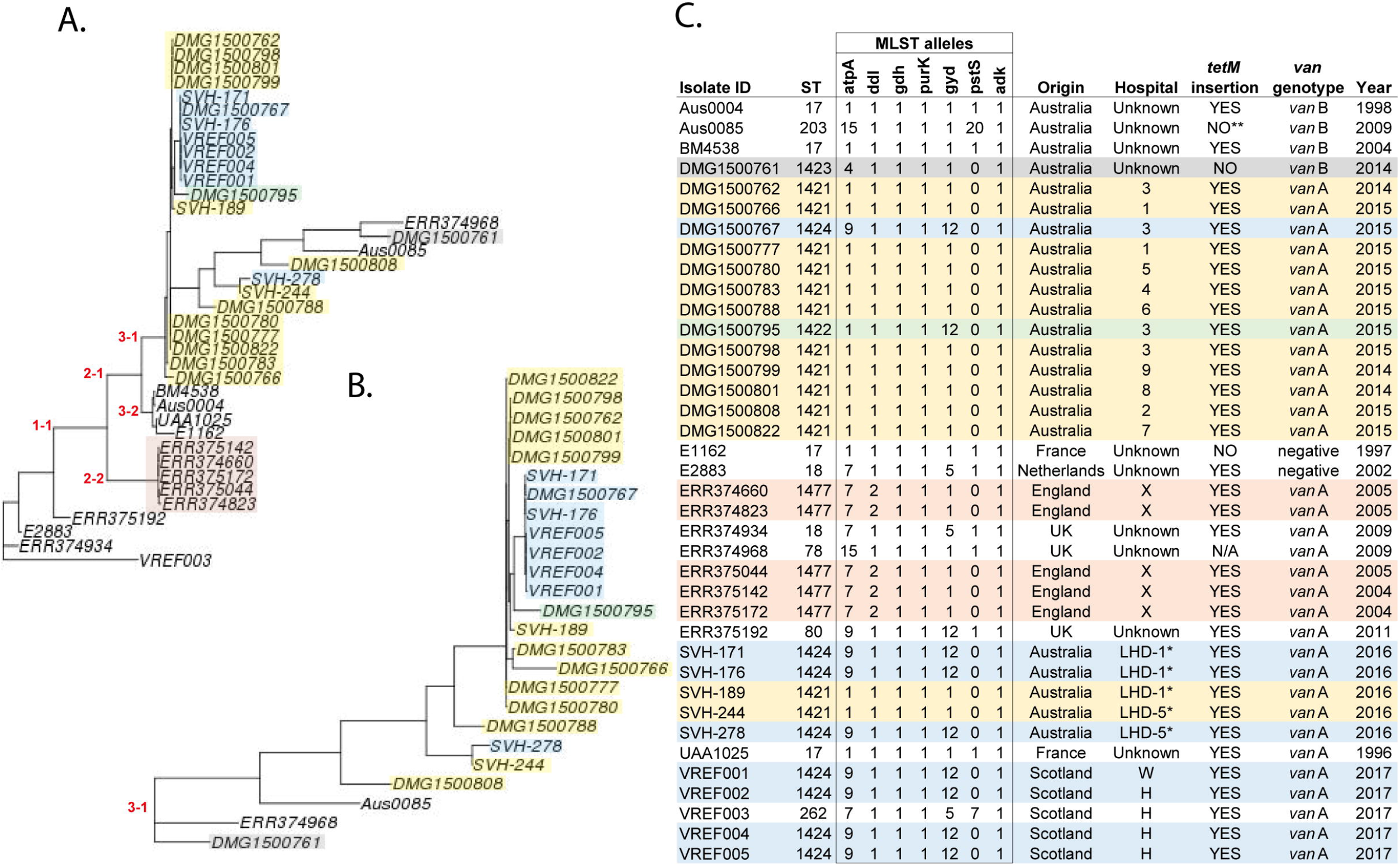
**A.** Phylogenetic tree of selected *E. faecium* genomes, constructed using the REALPHY (Reference sequence Alignment based Phylogeny builder) online tool, with Aus0004 and Aus0085 used as reference genomes. *pstS*-null genomes are present in subclades 2-2 (English isolates) and 3-1 (Australian and Scottish isolates), **B.** The phylogeny of strains clustering in subclade 3-1 (A) was further refined using REALPHY with Aus0085 and ERR374968 as references. **C.** The MLST sequence type (ST) and alleles, country of origin, *van* genotype, hospital and year of isolation are given for each sequenced genome. * Only the Local Health District (LHD) is known for Australian strains isolated in New South Wales hospitals during 2016; hospitals 3 and 9 are also in New South Wales jurisdiction, but their LHD is not known. Information is given on whether an insertion encompassing *dfrG* and two uncharacterised genes was found within *tetM* in these genomes. N/A: not applicable (no *tetM* sequences found). ** Although no such insertion was identified in Aus0085, there was a frameshift mutation within its *tetM* gene. *pstS*-null genomes of same ST are highlighted with the same colour.

## Discussion

MLST analysis is inferior to whole genome sequence analysis for outbreak investigations of VREfm [7], due to the high rate of recombination events occurring within the *E. faecium* chromosome that cannot be captured adequately by MLST analysis [27,28]. Furthermore, the emergence of non-typeable VREfm strains poses an extra obstacle in implementing the MLST scheme for VREfm phylogenetic analysis. Non-typeable VREfm strains have recently emerged and have been shown to be very rare in the UK: in a study encompassing whole genome sequencing of nearly 500 *E. faecium* healthcare-associated isolates from 2001-2011 in the UK and Ireland, there were only 5 cases of *pstS*-null VRE strains, all of which were isolated from a single English hospital, between 2004 and 2005 [7].The emergence of such strains was even more recent in Australia: the first two strains were isolated in 2013, but numbers increased rapidly to a total of 89 cases, by the end of 2015 [8,29,30] and to about 300 cases, by the end of 2016 [25,30,31].

Here, we report four additional non-typeable VREfm strains isolated in 2017, which are the first identified in Scotland. Their striking genomic similarity, combined with the fact that one was isolated from a different hospital, indicates that these, although deriving from a single strain type, are not necessarily hospital-acquired, but may be health-care associated, or spread within the community. Although transmission may have occurred through the frequent transfer of equipment, specimens, staff and patients between these two hospitals (located in the same health board just 14 miles apart), evidence suggests that VRE strains are also transmitted outside hospitals, such as in long term care facilities [6]. Additionally, VRE clones can disperse outside health-care facilities, since strains can be carried in patients for long periods, ranging from a few months to a few years following their discharge from hospital [32,33]. For the above reasons VRE infections may spread over long periods and across long distances. Indeed, our phylogenetic analysis is indicative of an inter-continental spread between Australia and Scotland: the closest phylogenetic relationships were identified among Australian and Scottish ST1424 clones; such 6-locus sequence type has been identified in 28.7% (58/202) of the recently sequenced *pstS*-null VREfm strains from New South Wales, Australia [25]. In addition, the spread of such strains in all health districts of New South Wales is also suggestive of inter-hospital transmission for this VREfm clone [25]. But even within the same hospital, VRE transmission routes can be prolonged and complex with closely related VRE strains re-appearing in the same ward after several months, or transmitted to patients located in different wards [34]. Indeed, we also observed re-appearance of a *pstS*-null strain (isolate VREF005) in the same hospital, seven months after discharge of the last carrier patient.

With the exception of one vancomycin-sensitive and one *vanB* type strain, all other (69) previously sequenced *pstS*-null *E. faecium* strains have been found to be *vanA* type [7,8], and consistent with these studies, the newly sequenced Scottish VREfm strains with the missing *pstS* locus were also *vanA* type. In fact, the vast majority (84.2%) of all *VanA E. faecium* 2016 isolates from New South Wales (Australia) sequenced, have been identified as *pstS*-null strains [25]; hence *pstS*-null VREfm appears to be a very successful *VanA* clone. We further identified that loss of *pstS* in *VanA* type VREfm is strongly associated with an insertion in a *tetM* gene (27/27 cases tested). Such insertion was absent only in the sole *vanB* type E. faecium strain of the *pstS*-null collection. As the 18 Australian *VanA* type *pstS*-null strains analysed here, were chosen to represent all 9 hospitals in the 3 health jurisdictions (or different local health district of the same jurisdiction) and phylogenetically distinct clades of the Australian *pstS*-null collection [8,25], it is highly likely that most (if not all) of the reported VanA-type *pstS*-null Australian strains excluded from our analysis (about 280 isolates) would contain a Tn5801-like transposon with an insertion in the *tetM* locus. The reason that these two features (missing *pstS* and *tetM* insertion) were co-selected in numerous VREs is currently unknown and deserves some further investigation. The *tetM* insertion leads to apparent inactivation of the resulting gene product towards tetracyclines, as can be inferred from Aus0004 [17], VREF001-2 and VREF004-5 (this study) susceptibility tests against tetracycline, doxycycline and minocycline. All sequenced isolates were resistant to trimethoprim, and the cause of such resistance is most likely the *dfrG* gene inserted into the *tetM* locus. None of the isolates sequenced had a tigecycline, Synercid, or linezolid resistant gene, and as a consequence they were all susceptible to these 3 drugs. Thus these antibiotics. which are frequently used for management of multi-drug-resistant enterococcal infections [1,35], may still be the favourable option for combating VREfm strains with a missing *pstS* locus.

The emergence of *pstS*-null VREfm strains in Scotland raises concerns for a potential outbreak: these strains were found to be highly similar to Australian *pstS*-null strains (this study), while the latter have very rapidly expanded to become the leading strain in Australia (2016), causing infections in numerous hospitals and in different health jurisdictions [8,25,26]. Furthermore, the missing *pstS* locus of such strains renders the - limited in resolution - MLST analysis even less suitable for VREfm outbreak investigations. Hence, our study, like previous ones [7,8,25], highlights the need for whole genome sequencing approaches in monitoring VREfm epidemiology.

## Acknowledgements

This work was supported by the Scottish Chief Scientist Office grant TCS/16/24 to NPT, SJD and ISH. We are very thankful to Gavin Paterson and Alison MacFadyen for reading the manuscript and for their useful comments.

